# Characterisation of *Pseudomonas aeruginosa’s* metal-responsive TonB-dependent transporters

**DOI:** 10.1101/2024.10.14.618150

**Authors:** Manon Ferry, Hortense Ferriz, Connor Sharp, Christos Paschalidis, Emmanuel Boutant, Isabelle J. Schalk, Olivier Cunrath

## Abstract

*Pseudomonas aeruginosa*, a versatile bacterium, relies on several TonB-dependent transporters (TBDTs) for nutrient acquisition (such as iron-siderophore complexes) and adaptation to various environments. While some TBDTs are well characterized, a significant number remain unexplored despite their potential role in pathogenicity. In this study, we developed fluorescent reporter plasmids to investigate TBDT promoter activity. Initially, we characterized their promoter activity in commonly used laboratory conditions, revealing diverse expression patterns among all TBDTs. Subsequently, we classified the TBDTs into distinct metal-responsive groups based on their stress-responsive behaviour, shedding light on their functional roles. Additionally, we show that these reporter constructs can be used as a powerful tool for siderophore detection. Finally, single cell analysis of TBDT promoter activity during coculture with an enterobactin-producing *Klebsiella pneumoniae* strain shows homogenous expression of key TBDT in *P. aeruginosa*. Our findings provide valuable insights into the expression profiles and functional diversity of *P. aeruginosa* TBDTs in complex conditions, reaching from commonly used lab media to complex co-culturing conditions.

## Introduction

*Pseudomonas aeruginosa* is a versatile and adaptable opportunistic pathogen known for causing severe infections in humans, particularly among immunocompromised individuals. To thrive and sustain its pathogenicity, *P. aeruginosa* requires an array of micronutrients, including essential metals such as iron and zinc. The uptake of these micronutrients, is mediated by metallophores—small organic molecules secreted by the bacterium to sequester metals with a very strong affinity from the environment. Notably, *P. aeruginosa* produces two siderophores (iron-chelator), pyoverdine and pyochelin, which vary in their iron-binding affinities and transport mechanisms (1–5). In addition, *P. aeruginosa* is also able to exploit exogenous siderophores produced by other microorganisms (5–16). Similarly, *P. aeruginosa* produces and uses the zincophore (zinc-chelators) pseudopalin for zinc acquisition (17). Once the metallophore-metal complex is formed in the extracellular space, it binds and is transported back into the bacterial cell via TonB-dependent transporters (TBDTs) – recently reviewed in (5). TBDTs are β-barrel proteins embedded in the outer membrane, crucial for the acquisition of micronutrients. Structurally, TBDTs form a conduit that facilitates the passage of these vital molecules into the periplasmic space. The function of TBDTs is closely associated with the inner membrane proteins TonB, ExbB and ExbD. Together, these proteins form a complex that utilizes the proton motive force across the inner membrane to energize the transport of substrates through the outer membrane via TBDTs (18).

In *P. aeruginosa*, the expression of metal-regulated TBDTs is tightly controlled by regulatory proteins such as the ferric uptake regulator (Fur) and the zinc uptake regulator (Zur) (19–22). These regulators repress TBDT expression in the presence of sufficient intracellular metal concentrations, in order to avoid metal intoxication. For a more precise regulation, in *P. aeruginosa* the expression of specific TBDTs is often induced by their corresponding ligands, further fine-tuning the nutrient acquisition process (5,7).

While the role of some TBDTs in metal acquisition—particularly involving siderophores for iron, zincophores for zinc, and other metal transport systems—has been investigated experimentally in *P. aeruginosa*, the functions of several other TBDTs remain still unknown. Despite their critical roles in pathogenicity and as potential gateways for siderophore-antibiotic conjugates, such as cefiderocol, a comprehensive understanding of all metal-responsive TBDTs in *P. aeruginosa* is still lacking. Furthermore, due to the absence of TBDTs expression profiles in commonly used lab media, a considerable uncertainty has arisen regarding the appropriate susceptibility testing for cefiderocol in Gram-negative bacteria (23,24).

This study characterizes the metal-responsive TBDTs in *P. aeruginosa*, shedding light on their structural and functional diversity. By elucidating the roles of some unknown TBDTs, we aime to help better understand their contributions to the pathogen’s physiology, which could pave the way for novel therapeutic strategies against *P. aeruginosa* infections.

### TonB-dependent transporters are weakly expressed in most laboratory conditions

In light of the growing antibiotic crisis and the crucial need for new gateways for antibiotics into the bacterial cell, we started to investigate the expression pattern of all 35 predicted TBDTs of the clinical isolate *P. aeruginosa* PAO1. Therefore, we constructed epitopic fluorescent transcriptional reporters using the promoter regions of each gene encoding a TBDT to drive the expression of YPet, a fluorescent protein that does not interfere with *P. aeruginosa’s* endogenous fluorescent siderophore pyoverdine (fig. S1A); combined to a constitutive expression of a second fluorescent protein, mCherry (fig. S1B).

Using this tool, we cultivated *P. aeruginosa* WT, each carrying one of the 35 TBDT reporter constructs in commonly used laboratory conditions, of which some are used to assess antibiotic susceptibility in clinic (25). We measured growth, mCherry and YPet fluorescence (Fig. 1 and fig. S2, S3). Total promoter activity was low for the vast majority of the transporters in all conditions, with activities observed for only a few transporters, namely the pyochelin siderophore transporter P*fptA*, the pyoverdine siderophore transporter P*fpvA* and the predicted cobalamin transporter P*btuB* (26) (Fig. 1A and fig. S2A). Importantly in Muller Hinton Broth (MHB), the culture medium that is most routinely used to assess antibiotic susceptibility (25), only minimal promoter activity was observed. Furthermore, the addition of sub-inhibitory concentrations of EDTA or apo-transferrin – both iron chelators – (fig. S4A, B) did not strongly affect promoter activity. Additionally, even the pre-treatment of the media using general metal-chelating resin Chelex^®^ only weakly changed promoter activity pattern. However, the addition of sub-inhibitory concentrations of calprotectin – a zinc-binding immune protein – (fig. S4C) increased the promoter activity of the zinc-responsive pseudopalin transporter P*cntO* as well as P*PA2911*, P*PA1922* (also named P*cirA*) and P*PA0781*, predicted to be involved in zinc uptake (17,27). Overall, general promoter activity was low in nutrient-rich growth conditions such as Lysogeny Broth (LB), Brain Heart Infusion medium (BHI) and Dulbecco’s Modified Eagle Medium (DMEM) supplemented with Fetal Bovine Serum (FBS). Addition of sub-inhibitory concentrations of EDTA (fig. S4D, E) only weakly alter promoter activity in LB or BHI. A strong increase in siderophore transporter activity was only observed when *P. aeruginosa* was cultured using defined or semi-defined media such as M9-succinate or Casamino Acid Medium (CAA), which are standard media for investigating iron starvation in *P. aeruginosa* (28) (Fig. 1A and fig. S2A).

**Figure 1.**
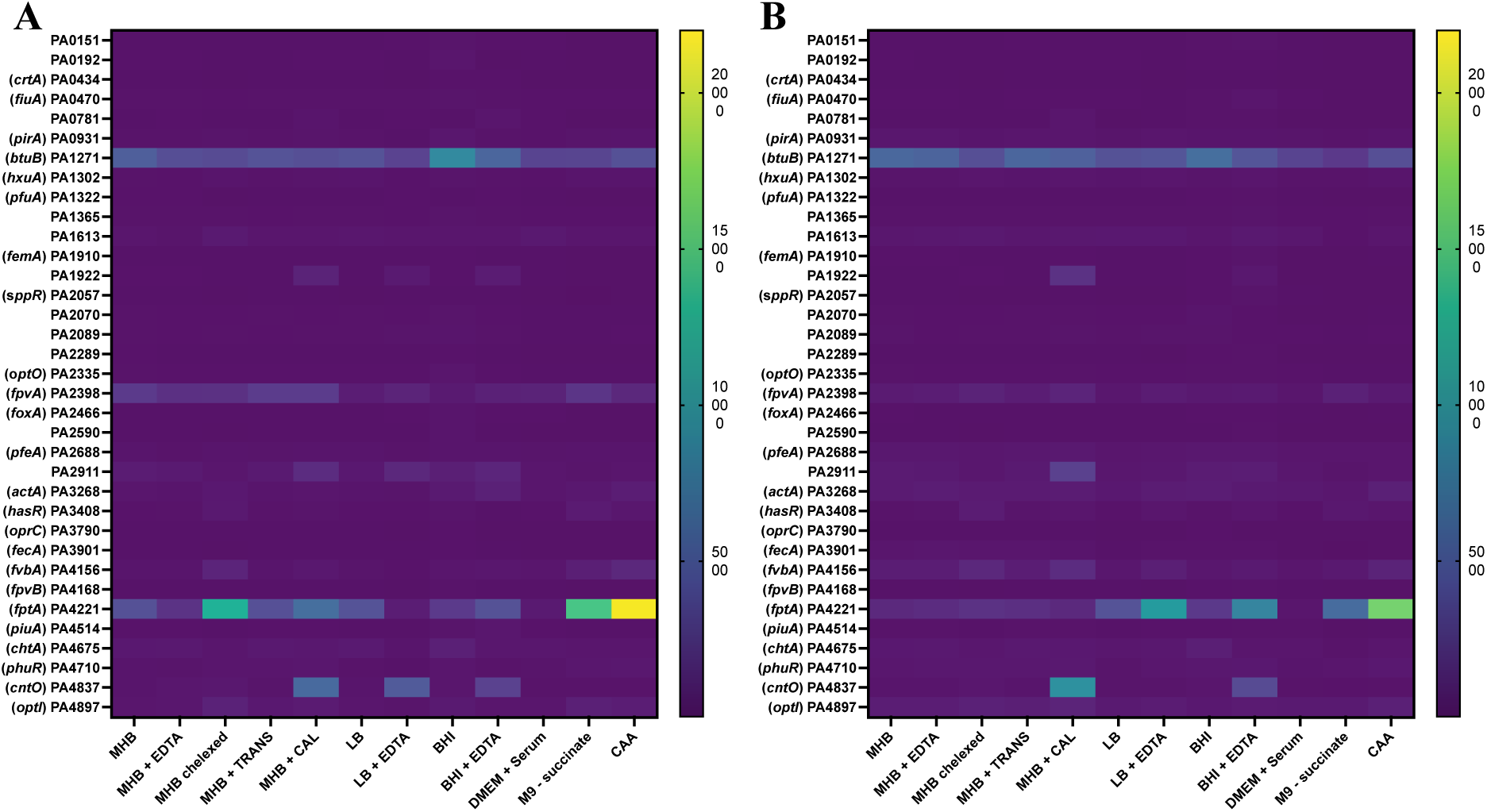
*P. aeruginosa* TBDT promoter activity in various growth condition. Absolute values of *P. aeruginosa* PAO1 WT (A) and its isogenic *pchA pvdF cntL* mutant (B) TBDT promoter activity measured by fluorescence (YPet) / Optical density at 600 nm. The colour code goes from dark blue (least expression) over green (medium expression) to yellow (higher expression). Bacteria carrying TBDT reporter plasmids were grown in media indicated on x-axis with supplementation of various additives at 37°C for 20 h, shaking. Muller Hinton Broth (MHB): + 300 µM of EDTA; Chelex^®^ treated; + 30 µg/ml of apo-transferrin (TRANS); + 30 µg/ml of calprotectin (CAL). LB: + 30 µM of EDTA. BHI: +100 µM of EDTA.

The ability of *P. aeruginosa* to produce its own metallophores has been shown to impact the expression of various exogenous siderophore transporters. As some clinical strains lose their ability to produce siderophores during chronic lung infection and switch towards different iron sources (29,30), we wanted to investigate the TBDT promoter activities in the absence of their endogenous metallophores. To evaluate the effect of endogenous metallophore production on TBDT’s promoter activity we generated an isogenic mutant lacking the biosynthesis of all three metallophores, namely pyochelin (*pchA*), pyoverdine (*pvdF*) and pseudopalin (*cntL*). *pchA pvdF cntL* deletion did not affect growth in rich conditions, but had a growth defect in the presence of metal (iron or zinc) restriction (fig. S4). We then tested if the absence of the endogenous metallophores had an effect on the TBDT promoter activities in using the same fluorescent reporters described above in the *pchA pvdF cntL* mutant strain. Generally, promoter activity was similar to WT pattern in all tested growth conditions. Interestingly, the absence of endogenous metallophore production slightly increased the metal starvation response in the presence of calprotectin and EDTA, but decreased in M9-succinate and CAA medium (Fig. 1B and fig. S2B, S3C, D, S4).

### Most TBDTs are responsive to metal starvation

To investigate the function of TBDTs in metal homeostasis, we cultivated our reporter strains in minimal media and exposed them to single stressor while monitoring their promoter activity. We used the semi-defined minimal medium CAA, which is routinely used for studying iron restriction. This medium has been shown to contain only around 20 nM of iron and induce endogenous iron uptake systems (28). Here, we could confirm that P*fptA* and P*fpvA*, both main endogenous siderophore transporters were highly upregulated in this condition (Fig. 1 and fig. S2). To investigate whether other TBDTs are involved in iron transport, we supplemented our medium with 10 µM FeCl_3_ and measured the promoter activity of all 35 transporters (Fig. 2A). It is well documented that the general iron uptake regulator Fur inhibits the transcription of the iron transporters at high cytoplasmic iron concentration (5,20). Interestingly, 18 different TBDTs were significantly repressed by the addition of iron, suggesting their involvement in iron acquisition. Of all 18 iron responsive transporters, three new transporters have been identified to be involved in iron acquisition, namely P*PA0151*, P*PA1365* and P*PA4897*. All 15 other TBDTs identified through this approach have been shown to be involved in iron acquisition, confirming the validity of our approach (Fig. 2A) (5).

**Figure 2.**
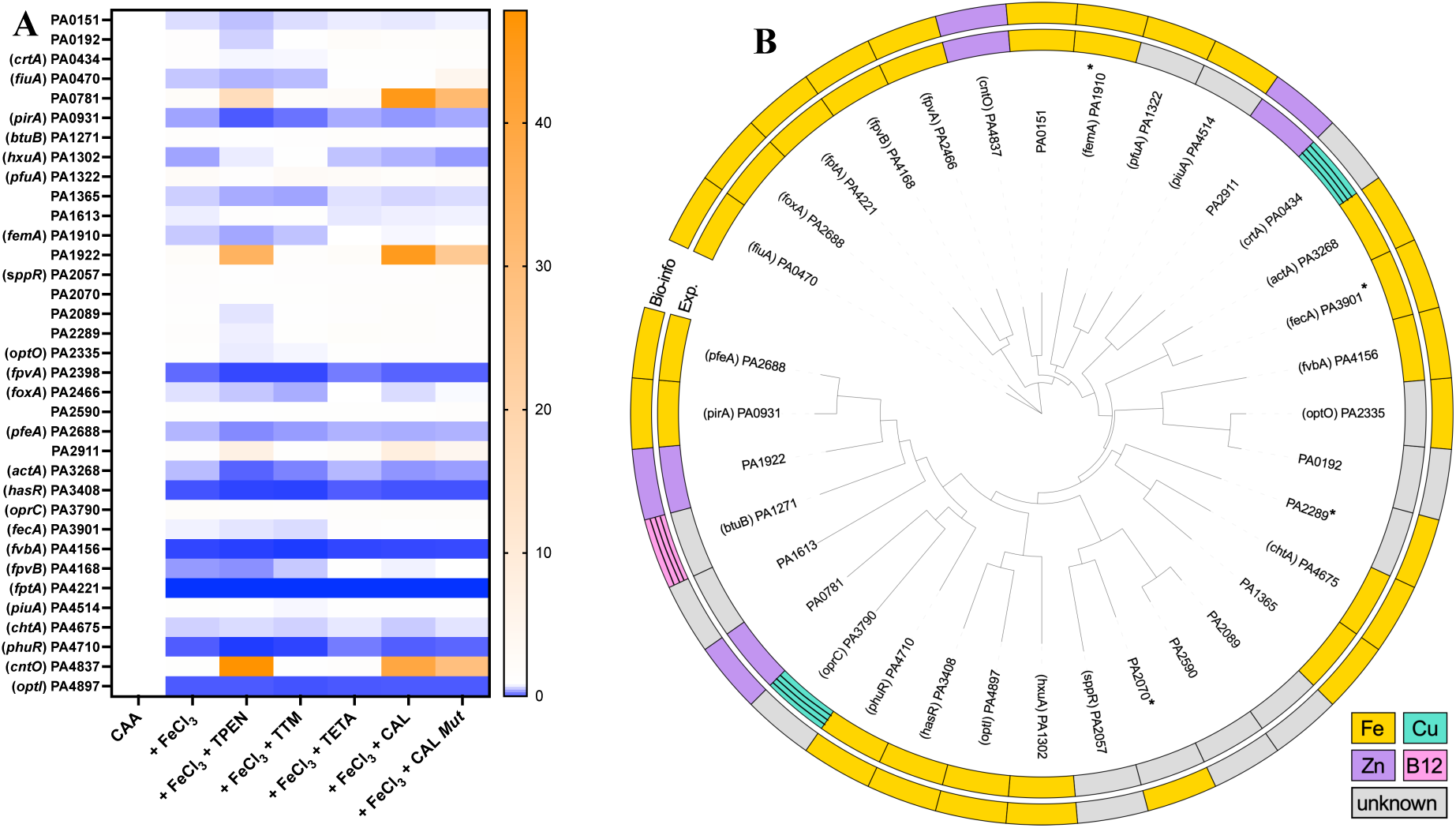
*P. aeruginosa* TBDTs are mostly metal responsive. (A) Fold change of TBDT promoter activity of *P. aeruginosa* PAO1 *pvdF pchA cntL* mutant, comparing promoter activity of strains grown in CAA in the absence and presence of single stressors. For each TBDT, the growth in CAA medium is the reference, and consequently the value of fold change is 1. Fold change of 1 was set to white, fold change lower than 1 is set to blue and fold change bigger than one is set to orange. Bacteria were grown at 37°C for 20 h, shaking supplemented with 10 µM of (FeCl_3_; TPEN; TTM, TETA) or 30µg/ml of (CAL, CAL *Mut*). (B) Phylogeny of 35 TBDT proteins. Full length protein sequences were aligned using clustalo and a phylogeny calculated using IQ-tree2. Inner ring indicates experimental prediction of import function. Outer ring shows the prediction from bioinformatic analysis of promoter regions. *Are indicated when Fur boxes were not found in the direct promoter region, but rather in the close genetic proximity. Hatched squares indicate experimental data or prediction from others (26,33,35).

To investigate if the remaining transporters were involved in metal acquisition other than iron, we first generated a zinc specific starvation growth condition. Therefore, the addition of the zinc chelator *N,N,N′,N′-*tetrakis(2-pyridinylmethyl)-1,2-ethanediamine (TPEN) in CAA which was supplemented with 10 µM FeCl_3_ allowed (i) the repression of iron responsive transporters and (ii) the upregulation of zinc responsive transporters (Fig. 2A and fig. S5). This helped us to identify four zinc responsive TBDTs (P*PA0781*, P*PA1922*, P*PA2911* and P*cntO*), confirming previous findings (17,27). Additionally, our approach suggests that all other TBDTs may not be involved in zinc uptake.

The same approach was conducted to generate a copper specific starvation, using either tetrathiomolybdate (TTM) or triethylenetetramine (TETA) known to be strong copper chelators. Unfortunately, none of those condition resulted in a measurable increase of promoter activity (Fig. 2A). Even after the deletion of the general copper homeostasis regulator gene *cueR*, which should result in an increased transcription of genes involved in copper uptake (31,32), no detectable copper response was measured for any TBDT, including the two described copper responsive transporters P*oprC* and P*ctrA* (fig. S6) (33–35).

Finally, to verify if some TBDTs are responsive to other transition metals such as manganese and due to the extreme challenge of having a manganese specific starvation response, we turned to the use of the metal binding protein calprotectin. Calprotectin is a metal-binding protein complex, involved in the innate nutritional immune response (36,37). Natural calprotectin contains two transition metal binding sites: H3N (binding Mn, Cu) and H6 (binding Zn, Fe, Ni) (36). Using sub-inhibitory concentration of calprotectin (fig. S7) in our experimental conditions we could monitor a zinc-starvation response, similar to the one induced by TPEN. Additionally, the use of the mutated calprotectin variant with alterations in the H3N site with substitutions H103N, H104N, and H105N, which disrupt its metal binding capability resulting in the abolishment of manganese and copper binding, did not show a different promoter activity pattern either (Fig. 2A). This finding suggests that none of the above-mentioned transporters are involved in the uptake of manganese. Finally, even in those condition, no copper starvation response was observed.

### Sequences can only partially predict transporter function

Protein function can often be predicted from sequence. Here we used sequence data from all 35 TBDT of *P. aeruginosa* combined with our experimental predictions of function to examine if protein sequence can predict the functional role of TBDTs. Using full length protein sequences of 35 TBDTs, we did not find an association between sequence homology and TBDT function (Fig. 2B). TBDTs involved in the uptake of zinc and iron were spread throughout a phylogeny of TBDT sequences, and individual TBDT involved in zinc uptake were closely related to those involved in the uptake of iron, suggesting that full length sequence is not predictive of functional role. Using alignments of TBDT functional domains, (Plug: PF07715 and TonB_dep_Rep: PF00593) did not improve the ability to predict its functional role (fig. S8). While protein sequence was not predictive of a transporter’s function, we sought to test if the promoter regions might be more predictive. Using MAST (from the MEME Suite) we were able to identify either Fur or Zur boxes in the promoter regions of many TBDTs, which largely agreed with experimental predictions of TBDT function (Fig. 2B and fig. S8). Interestingly, FecA and FemA, known to be involved in Fe acquisition through the iron chelators citrate (12) and mycobactin (7) respectively, did not have a detectable Fur box in their promoter region. While our experimental results and previously published data show clear iron regulation, we sought to search for Fur boxes in the close genetic regions close to TBDTs. Using this approach, we could find four TBDTs (namely FemA, FecA, PA2070 and PA2289) which had predicted Fur boxes in their operons (Fig 2B TBDT with * and fig. S8). Those transporters may not directly be regulated via Fur, but Fur may rather bind on the promoter region of their positive regulators such as FecI and FemI, respectively, explaining our and other’s previous results (5,7). Combining our experimental approach with this bioinformatic analysis, we suggest the involvement of the three new unidentified TBDTs in the acquisition of iron (PA0151, PA1365 and PA4897) (Fig. 2). Additionally, three transporters (namely PA1322, PA2335 and PA4514), with potential Fur boxes in their promoter regions and two additional transporters with Fur boxes in their close genetic environment (PA2070 and PA2289) have been identified bioinformatically only. Our experimental data could not reveal any significant promoter activity in our growth conditions, which might be due to very low promoter activity resulting in undetectable variation or simply due to false positives during the bioinformatic approach.

### Reporter constructs can be used for siderophore detection

Most TBDTs involved in exogenous siderophore uptake are highly upregulated in the presence of their respective ligand, especially in a strain that does not produce endogenous siderophores. This upregulation is mediated via Sigma/Anti-sigma factors, two component systems or cytoplasmic AraC-regulators and IclR-regulators, leading to a specific increased expression of their respective transporter (5). Citrate for example is transported through the TBDT FecA (12). The transport of citrate activates the sigma factor FecI via the inner membrane anti-sigma factor FecR, leading to the upregulation of FecA (7). The same is true for ferrichrome, which leads to an upregulation of its TBDT FiuA (9) via FiuI/R (7), for ferrioxamine-E and its transporter FoxA (38) via FoxI/R (7) and enterobactin for PfeA (6) via PfeS/R (39). This specific increased promoter activity may be used as a detection strategy for detecting specific molecules in complex media. To test this, we choose various siderophores known to induce the expression of their respective transporters. All tested siderophore-transporter combinations were tested on a *P. aeruginosa* PAO1 *pvdF pchA cntL* mutant strain and showed a specific increase in promoter activity, while not influencing final bacterial abundance (citrate – FecA; ferrichrome – FiuA; ferrioxamine-E – FoxA; enterobactin – PfeA) (Fig. 3A and fig. S9). In addition to known tested siderophores, we also tested spermidine, a siderophore precursor which has been suggested to be able to play a role in iron homeostasis (40). No upregulation of any transporter has been measured. Finally, for all tested molecules, a strong down-regulation of the endogenous-siderophore transporter FptA was observed, which is in accordance with published data (5,41) (Fig. 3A).

**Figure 3.**
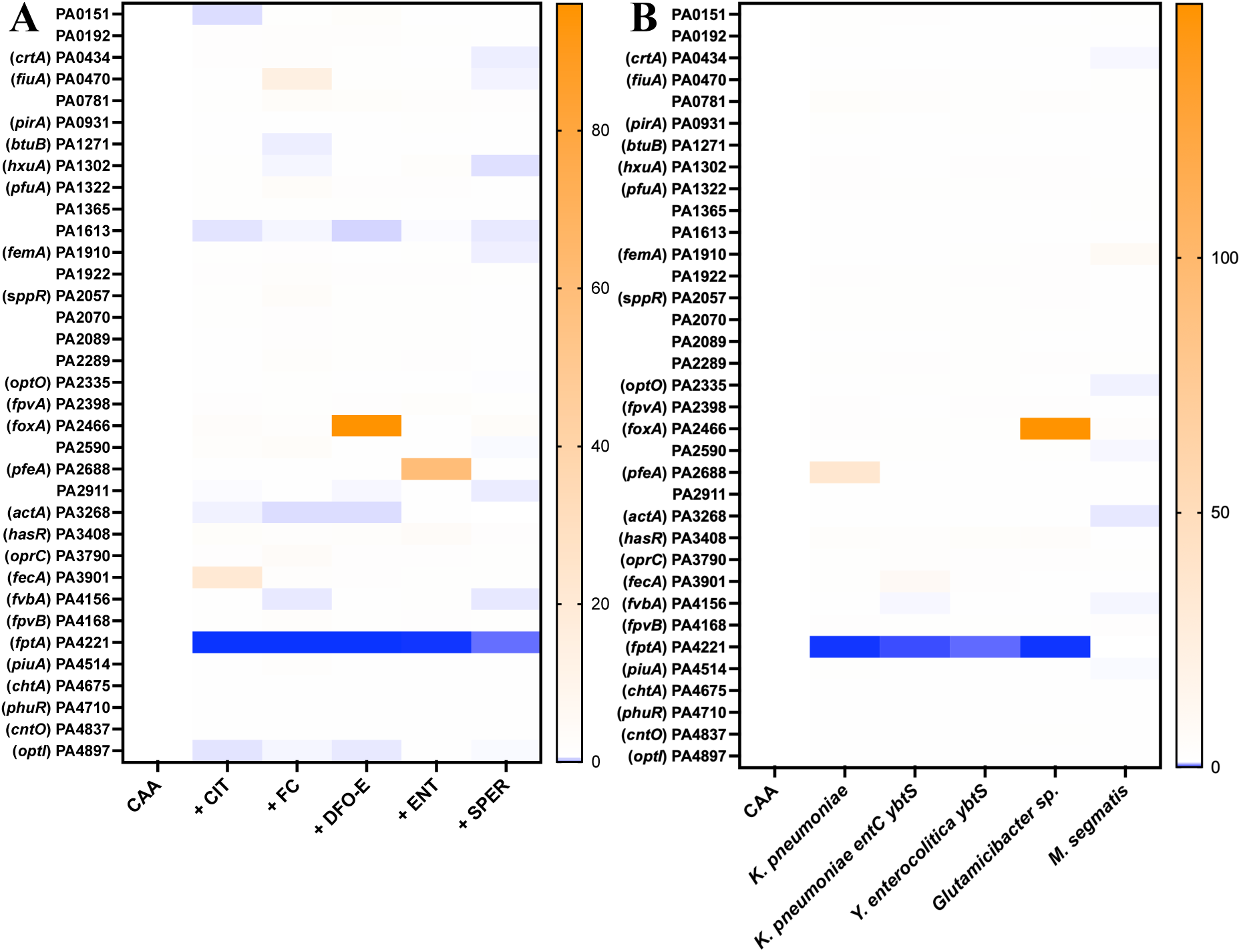
Reporter constructs can be used for siderophore detection. (A) Fold change of TBDT promoter activity of *P. aeruginosa* PAO1 *pvdF pchA cntL* mutant, comparing promoter activity of strains grown in CAA in the absence and presence of single molecules. For each TBDT, the growth in CAA medium is the reference, and consequently the value of fold change is 1. Fold change of 1 was set to white, fold change lower than 1 is set to blue and fold change bigger than one is set to orange. Bacteria were grown at 37°C for 20 h, shaking supplemented with 1 mM of CIT – citrate, or 10 µM of FC - ferrichrome; DFO-E – desferroxamine E; ENT – enterobactin; SPER – spermidine. (B) Fold change of TBDT promoter activity grown in CAA medium exposed to bacterial supernatants.

We then set out to test if our reporter constructs would also detect these siderophores in complex growth environments. We therefore used sterile filtered bacterial supernatant from known siderophore producers. We used the supernatant of *Klebsiella pneumoniae* subsp. *pneumoniae* (DSM 30104), possessing enterobactin and yersiniabactin biosynthesis machineries and its isogenic mutant lacking key genes in enterobactin and yersiniabactin production *entC ybtS*. Using our reporter construct and a *K. pneumoniae* WT supernatant, we could detect a strong increase in P*pfeA* activity and decreased P*fptA* activity, in line with *P. aeruginosa’s* exploitation strategy due to the presence of enterobactin in the supernatant of the wild type strain (Fig. 3B and fig. S10). On the contrary, no P*pfeA* upregulation was detected in the double mutant *K. pneumoniae entC ybtS*, which had no detectable siderophore production in the supernatant anymore, as shown using the Chromazurol S (CAS) assay (fig. S11). Surprisingly, a weak increase of the citrate promoter P*fecA* was measured when *P. aeruginosa* was exposed to a *K. pneumoniae entC ybtS* supernatant.

Interestingly, no TBDT other than enterobactin transporter P*pfeA* was upregulated in the presence of *K. pneumoniae* WT supernatant, despite it being reported that yersiniabactin would induce P*femA* (42). This absence of induction could be either due to siderophore competition, where all iron ions would be complexed by enterobactin (which has a stronger affinity for iron than yersiniabactin); or due the absence of yersiniabactin expression. While it has been reported that some *Klebsiella* strains do not produce yersiniabactin in large amounts (43) we confirmed this hypothesis showing that the single mutant *entC* did not show any detectable siderophore production using the CAS assay (fig. S11). To further understand if yersiniabactin might induce P*femA*, we turned to another pathogen known to produce high amounts of yersiniabactin. We therefore tested the supernatant of *Yersinia enterocolitica* subsp. *enterocolitica* (DSM 27689) and its isogenic mutant *Y. enterocolitica ybtS* lacking a key gene for yersiniabactin biosynthesis. While both *Y. enterocolitica* strains grew similarly in the tested iron-deprived medium, only the *Y. enterocolitica* wild-type strain demonstrated significant siderophore production using the CAS assay (fig. S11). Surprisingly, using the supernatant of the *Y. enterocolitica* wild-type strain, the growth of our *P. aeruginosa* reporter strains was fully repressed (fig. S10C, D). This repression was abolished when using the supernatant of the *Y. enterocolitica ybtS* mutant strain. Using these data, we suggest that *P. aeruginosa* might actually not be able to use yersiniabactin, contradicting previous results (42). Finally, using the supernatant of an environmental bacterial ferrioxamine-producing *Glutamicibacter sp.* strain and a mycobactin-producing *Mycobacterium smegmatis* strain (DSM 43756), we could measure increased P*foxA* and P*femA* promoter activity, respectively (Fig. 3B). Taken together, we could show that the above described reporter constructs could also be used as a rapid and efficient siderophore detection tool in complex growth conditions.

### Enterobactin exploitation is homogenous in co-culture

Single cell measurements often differ significantly from whole population averages, masking critical biological insights (44). Our reporter constructs facilitate single cell analysis via flow cytometry, enabling precise quantification and characterization of individual cell variations within a population. The constitutive expression of mCherry by all *P. aeruginosa* cells allowed proper separation of *P. aeruginosa* cells from background noise and allowed single cell analysis of promoter activity (fig. S12). Single cell analysis of *P. aeruginosa’s* major TBDT promoter activities in our laboratory conditions showed homogenous expression of the predicted cobalamin transporter P*btuB*, the endogenous zincophore pseudopaline transporter P*cntO*, the two endogenous siderophore pyoverdine transporters P*fpvA* and P*fpvB* and the endogenous siderophore pyochelin transporters P*fptA* (Fig. 4). Additionally, we explored promoter activity of the gene encoding for enterobactin transporter P*pfeA* during co-culture with our enterobactin-producing *K. pneumoniae* strain or its isogenic mutant *entC ybtS*. Here, we grew *K. pneumoniae* WT strain (which was carrying a plasmid with constitutive blue fluorescence for clean separation – fig. S12) together with *P. aeruginosa* and we could show that during co-culture, *P. aeruginosa*’s promoter activity of the gene-encoding enterobactin transporter P*pfeA* was strongly homogenously upregulated, suggesting that *K. pneumoniae* is producing significant amounts of enterobactin when competing with *P. aeruginosa* and that the latter successfully exploits this siderophore. This observation was abolished when co-culturing with a *K. pneumoniae entC ybtS* strain, which is unable to produce enterobactin. This upregulation was homogenous within the whole population of *P. aeruginosa* (Fig. 4). Interestingly, no down-regulation of the endogenous pyochelin pathway was observed in this condition.

**Figure 4.**
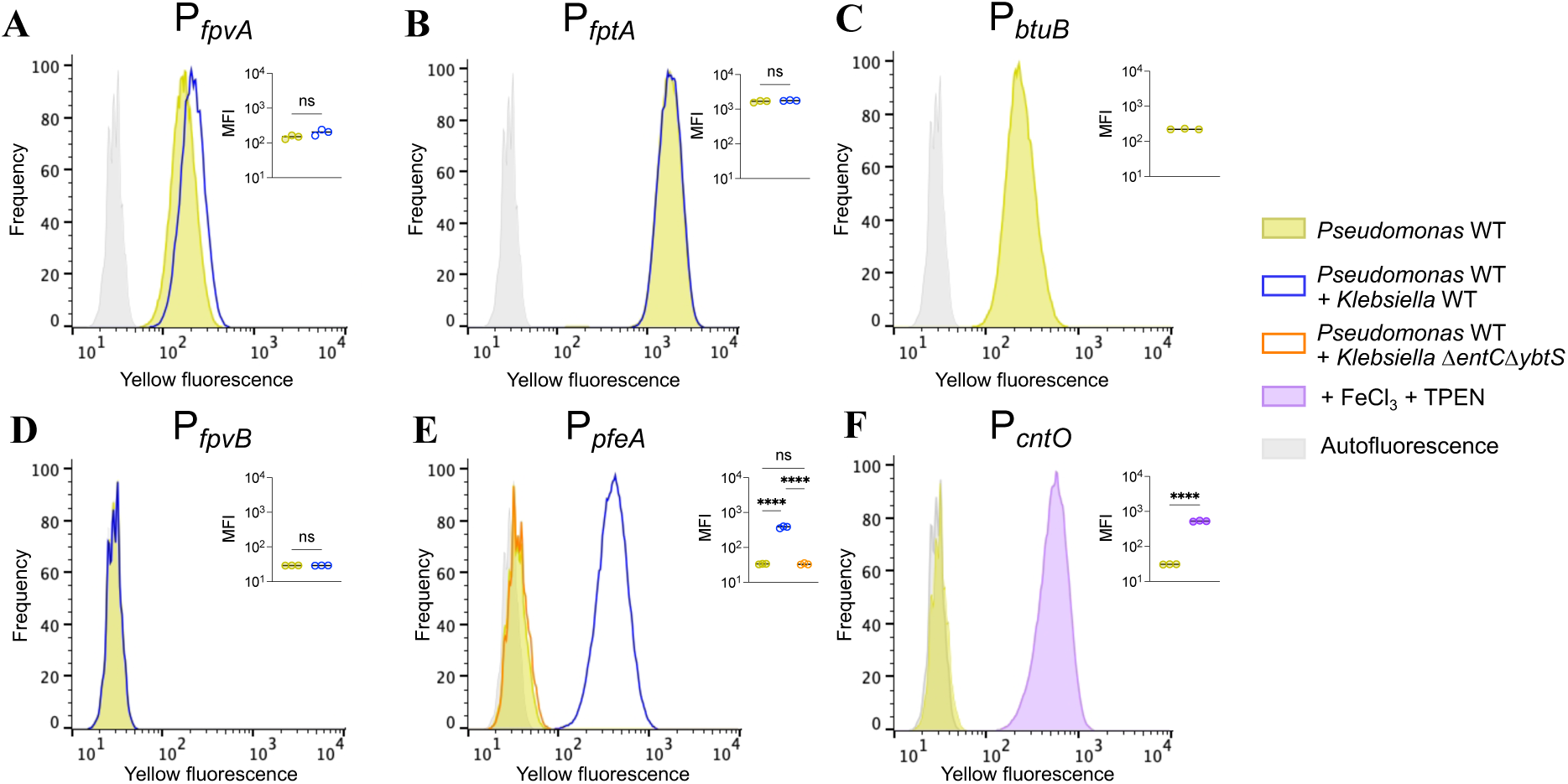
TBDT expression is homogenous in laboratory conditions and during co-culture. Panels show the yellow fluorescence of *Pseudomonas* strains expressing YPet for key TBDTs genes under different conditions (A – pyoverdine transporter FpvA, B – pyochelin transporter FptA, C – cobalamin transporter BtuB, D – hydroxamate transporter FpvB, E – enterobactin transporter PfeA and F – pseudopaline transporter CntO). Filled histograms represent monocultures of *Pseudomonas* strains during iron starvation (CAA) (yellow histograms) or zinc starvation (CAA + 10 µM FeCl_3_ + 10 µM TPEN) (purple histogram) conditions. Empty histograms represent the yellow fluorescence of *Pseudomonas* in co-cultures with *Klebsiella* strains that produce (WT) (blue histograms) or not (*entC ybtS*) (orange histogram) siderophores. Gray areas represent autofluorescence. MFI indicates median fluorescence intensity of YPet in mCherry subsets (for the gating strategy used see Fig. S12). Statistical significance was accessed using Mann-Whitney tests with a significance threshold of P <0.05.

## Discussion

TBDTs are essential for efficient energy-dependent transport of complex compounds that are unable to diffuse through porins across the outer membrane of Gram-negative bacteria. Our comprehensive investigation of all 35 TBDT promoter activities in a clinical *P. aeruginosa* isolate combined with our bioinformatic approach provided a unique opportunity for determining metal-dependent TBDT function. First, we show that most transporters are not or only weakly expressed in laboratory growth condition. This is especially true for Mueller Hinton Broth, which is still largely used for assessing antibiotic susceptibility, possibly masking key physiological features of *P. aeruginosa*’s membrane permeability. Our results provide additional weight to the idea that the use of a more defined medium might be better suited for drug susceptibility, especially for drugs that may enter the bacterial cell via TBDTs. Further investigations are crucially needed to find a more clinically relevant medium for assessing antibiotic susceptibility.

Additionally, combining our experimental and bioinformatic approach, we show for the first time the involvement of 18 TBDTs in iron homeostasis, of which three are newly identified transporters. These transporters might be involved in the uptake of iron complexes such as exogenous siderophores that *P. aeruginosa* might encounter during its versatile life-style. In addition, five other TBDTs were bioinformatically predicted to be iron regulated, which haven’t been confirmed experimentally. We also show that only four out of the 35 TBDTs are involved in zinc uptake. No manganese or copper response was measured for any TBDT, which may come from our experimental detection threshold. To conclude, we show that at least 18 TBDTs are involved in iron acquisition and four TBDTs in zinc uptake, one TBDT may be involved in cobalamin uptake (which we do not formally assess here), two TBDTs may be responsive to copper starvation (though our data could not confirm these results) and 11 TBDTs may actually not be involved in metal uptake, but rather involved in the uptake of other metabolites or complex carbohydrates.

Subsequently, we also show that our reporters can be used as a powerful tool allowing rapid and accurate detection of various siderophores in complex media. With this, we could show that *P. aeruginosa* preferentially uses enterobactin when grown in an enterobactin containing *K. pneumoniae* supernatant, but that when exposed to a *K. pneumoniae* supernatant lacking enterobactin production, secondary transporters, such as FecA, may be of important use. We also suggest that *P. aeruginosa* is unable to exploit yersiniabactin, as the supernatant containing yersiniabactin is inhibitory for a *P. aeruginosa* strain unable to produce its own siderophore, while a *Y. enterocolitica* supernatant lacking yersiniabactin is not inhibitory. The controversary result may be due to the instability of yersiniabactin which as a purified molecule may rapidly break down and explain the previous published phenotype (42). Finally, single cell analysis shows that pyochelin, pyoverdine and pseudopalin acquisition, cobalamin uptake and enterobactin exploitation depict homogenous expression pattern in our mono- and co-culturing conditions. These results shed light into the versatility of *P. aeruginosa’s* metal uptake mechanisms and its phenotypic adaptation to complex growth conditions. Future research should further characterize the remaining metal none-responsive TBDTs with no known function which will lead us to a more complete understanding of *P. aeruginosa’s* versality and adaptability to complex and varying environments.

## Acknowledgments

We are indebted to members of the MMBCA lab for discussion. Thank you to Walter Chazin for providing WT and mutant calprotectin. Thank you to Kieran Bates for providing the *Glutamicibacter* strain.

## Funding

MF and CP were supported by an MRT Studentship. This work was supported by IdEx 2022 « Attractivité » University of Strasbourg and by a grant from the Agence Nationale de la Recherche (ANR, grant number: ANR-22- CE44-0024-01, acronyme: IRUPP). We also acknowledge the Interdisciplinary Thematic Institute (ITI) InnoVec (Innovative Vectorization of Biomolecules, IdEx, ANR-10-IDEX- 0002).

## Author contributions

Conceptualisation: OC. Methodology: MF, CS and OC. Investigation: MF, HF, CS, CP, EB, OC. Visualisation: MF, CS and OC. Funding acquisition: IJS, OC. Supervision: OC. Writing – original draft: OC. Writing – review and editing: MF, CP, IJS and OC.

## Competing interests

None.

## Data and materials availability

All data are available in the main text or supplementary materials.

## Methods

### Media

Rich media described in the main text were prepared according to manufactures’ recommendation. Lysogeny broth (**LB**) (AthenaES Ref:0102); BBL^TM^ Mueller Hinton II broth, Cation Adjusted (**MHB**) (BD Ref: 212322); Bacto^TM^ Brain Heart Infusion (**BHI**) (BD Ref: 237500); Dulbecco’s Modified Eagle Medium (**DMEM**) (Gibco^TM^ Ref:41965-039); heat-treated Fetal Bovine Serum (**FBS**) (Dutscher Ref: S1810-500). Minimal media were prepared as follows. Casamino Acid Medium (**CAA**) – 5 g/L Bacto^TM^ Casamino acid (BD Ref: 223050), 1.46 g/L K_2_HPO_4_ 3H_2_O (Carlo Erba Ref: 471767), 0.25 g/L MgSO_4_ 7H_2_O (Merck Ref: 1.05886.1000); Succinate medium (**SM**) – 6 g/L K_2_HPO_4_ 3H_2_O (Carlo Erba Ref: 471767), 3 g/L KH_2_PO_4_ 3H_2_O (Carlo Erba Ref:361507), 1 g/L [NH_4_]_2_SO_4_ (Merck Ref: 1.01217.1000), 0.2 g/L MgSO_4_ 7H_2_O (Merck Ref: 1.05886.1000), 4 g/L succinic acid (Sigma-Aldrich Ref: S3674) and pH was adjusted to 7.0 by using NaOH. Low phosphate M9 (**lpM9**) – 1.28 g/l Na_2_HPO_4_ (Merck Ref: 231-448-7), 0.3 g/l KH_2_PO_4_ (VWR Ref: 26936.260), 0.5 g/l NaCl (Euromedex Ref: 1112-C), 1 g/l NH_4_Cl (Merck Ref: 1145) and 10 ml/l Goodiemix. <u>Goodiemix solution</u> consists of 94.89 g/L MgSO_4_ 7H_2_O (Merck Ref: 1.05886.1000), 1.11 g/L CaCl_2_ (Sigma-Aldrich Ref: 31307), 0.033727 g/L thiamine hydrochloride (Sigma-Aldrich Ref: T4625-10G) and 125 ml/l trace element solution [13.08 g/l MgCl_2_ 6H_2_O (Merck Ref: 1.05833.1000), 2 g/l CaCl_2_ (Sigma-Aldrich Ref: 31307), 0.90 g/l ZnSO_4_ (Strem chemicals Ref: 93-3045), 0.85 g/l MnSO_4_ · H_2_O (Bio Basic Ref: MB0334), 0.24 g/l CuSO_4_ · 5 H_2_O (Strem Ref: 50-901-14907), 0.06 g/l H_3_BO_3_ (Sigma-Aldrich Ref: B0394-500G), 51 ml/l 1M HCl (Thermo-Fischer Ref: H/1200/PB15)]. Glucose (Euromedex Ref: UG3050) was used as carbon source (4 g/l final concentration).

### Bacterial strains

A full list of bacterial strains used in this study is provided in Table S1. *Pseudomonas aeruginosa* PAO1 (DSMZ 22644) was used in all experiments. *P. aeruginosa* PAO1 WT, *K. pneumoniae* WT and mutant strains were routinely streaked on LB agar plates containing appropriate antibiotics and incubated at 37°C for 16 h. Single colonies were picked and grown in CAA supplemented with 1µM FeCl_3_ and appropriate antibiotics (Table S1) and incubated at 37°C, shaking at 200 rpm for 16h. *Escherichia coli* was routinely grown in LB containing appropriate antibiotics and incubated at 37°C, shaking at 200 rpm for 16 h.

### Genetic engineering of bacterial strains

Gene deletions were generated as described in (45). Briefly, 700 base pairs upstream and downstream of region to be deleted were PCR amplified (Phusion^®^, NEB) and inserted into the suicide vector pEXG2, which was linearized using the restriction enzymes BamHI and EcoRI, using the NEBuilder^®^ HiFi DNA Assembly Master Mix (NEB #E2621L). The plasmid was introduced into an *E. coli* strain (TOP10). After 6h of mating between the plasmid-containing donor *E. coli* strain, the *E. coli* helper strain and the recipient *P. aeruginosa* strain, trans-conjugants were selected on LB plates containing 30 µg/mL gentamicin and 10 µg/mL chloramphenicol. Counter-selection was performed on no-salt LB plates supplemented with 10% of sucrose at 30°C. Mutants were screened by colony PCR using described primers and sequence was verified using Sanger sequencing (Eurofins). Primers used for genetic engineering in this study are listed in Table S2.

### Genetic engineering of reporter plasmids

All plasmids used in this study are listed in Table S3. Reporter plasmids were designed using a RK2 *oriV* backbone with a gentamicin resistance cassette to which P*_pilM_*-*mCherry* and P*_foxA_*-*ypet* were inserted. This plasmid was linearized using primers oOPC-1028 and oOPC-1029. Promoter region were amplified using primers described in Table S2. Promoter region were amplified by selecting the entire inter-genetic region and the first 45bp (15 codons) of the gene of interest and inserted *in frame* before a 32bp-long RBS containing region prior to the *ypet* gene. Promoter fragment and linearized vector were ligated using NEBuilder Assembly kit (E5520S – New England Biolabs). Ligated plasmids were transformed into heat-shock competent TOP10 *E. coli* cells. Plasmids were sequenced using Sanger sequencing and electroporated into recipient *P. aeruginosa* strains. For the electroporation *P. aeruginosa* was grown overnight in LB at 37°C. 1 ml of *P. aeruginosa* culture was washed three times at room temperature using a 300 mM sucrose solution. Final pellet was resuspended in 100 µl of 300 mM sucrose solution and electroporated with 100 ng of purified plasmid followed by immediate supplementation of 900 µl Super Optimal broth with Catabolite repression (SOC). Cells were incubated for 1h at 37°C and plated on LB plates containing 30 µg/ml of gentamicin and incubated at 37°C overnight.

### Bacterial growth kinetics

*P. aeruginosa* PAO1 WT or mutant strains were precultured as described above. Overnight cultures were harvested and OD_600 nm_ was measured using an Eppendorf BioPhotometer^TM^. OD_600 nm_ was adjusted to final OD_600 nm_ of 0.01. 96 well plates (Greiner Bio-One #655161) were prepared containing the described medium with various concentrations of chemical supplements. All chemical supplements and their respective solubilizing agent are listed in Table S4. 96 well plates were incubated in a Tecan plate reader (Series Infinite M200PRO), measuring OD_600 nm_ every 15 minutes for 24 h. Data were plotted using GraphPad Prism 10.

### Fluorescence measurement

*P. aeruginosa* carrying a low copy number plasmid with various P_TBDT_-*ypet* were precultured in CAA supplemented with 1 µM FeCl_3_ and 30 µg/mL gentamicin overnight at 37°C, shaking at 200 rpm. Overnight cultures were diluted 1/10 in a 96-well microplate in CAA supplemented with 30 µg/mL gentamicin. This 1/10 dilution was used to inoculate freshly prepared microplates, which contained the growth conditions described in main text. Plates were incubated for 20 h at 37°C, shaking at 600 rpm. After 20 h incubation, the microplates were centrifuged using a microplate centrifuge (Sigma Laborzentrifugen^TM^ 10155) at 3500 rpm for 10 min at room temperature. Supernatant were removed and bacterial pellets were resuspended in PBS. Optical density (600 nm), Ypet fluorescence (ex: 500 nm, em:540 nm) and mCherry fluorescence (ex: 570 nm, em: 610 nm) were measured using a TECAN plate reader (Series Infinite M200PRO). Data were plotted using GraphPad Prism 10.

### Co-cultivation with *Klebsiella pneumoniae*

From a single colony, *P. aeruginosa* carrying the reporter plasmid pOPC-249 (P*_pfeA_*-*yept*-P*_pilM_*-*mcherry*) and *K. pneumoniae* WT or *entC ybtS* mutant strains carrying pBC-22 (P*_ybaJ_*-*mtagbfp2*) are independently cultured for 20 hours in CAA medium (pH 7.4) supplemented with 1 µM FeCl_3_ and the appropriate antibiotics. For the co-cultivation, 1 ml of each culture is washed twice with fresh CAA medium, and 100 µL of each adjusted suspension to an OD_600 nm_ of 0.1, are then mixed in a tube containing 1800 µL of fresh CAA medium plus appropriate antibiotics. Consequently, the final OD_600 nm_ in the co-culture is 0.01 (0.005 per strains). The cultures are incubated at 37°C, shaking at 220 rpm for 24 hours.

### Flow Cytometry

For each culture tube, 1 µl was fixed in 1 ml of ice-cold PBS containing 4% PFA for 2 h on ice. All samples were kept on ice until and during analysis. Relevant spectral parameters were recorded in a FACS equipped with 405, 488, 561 and 637 nm lasers (BD FACSymphony A1 Cell Analyzer), using minimal thresholds on SSC and FSC to exclude electronic noise. We used the following channels: YPet, ex: 488 nm, em: 530/30 nm 505 nm LP (“yellow”); mCHERRY, ex: 561 nm, em: 610/20 nm 595 nm LP (“red”); BFP, ex: 405 nm, em: 450/50 nm 410 nmLP (“blue”).

### Chrome azurol S

CAS solution was prepared as described by Schwyn and Neilands (46). All reagents (Chrome Azurol S, HDTMA, anhydrous piperazine and 5-sulfosalicylic acid) were purchased from Sigma-Aldrich. Bacterial supernatants were filtered-sterilized on a 0,22 µm membrane. 100 µL of the filtered supernatant was mixed with 100 µL of CAS solution in a well of a 96-well microplate with flat bottoms (Greiner Bio-One #655161). Controls included uninoculated media mixed with CAS solution and an iron-free variant of the CAS solution (APO-CAS). The plate was then incubated in a dark environment at room temperature for 1 hour and 30 minutes. The optical density at 630 nm (OD_630 nm_) was measured using a TECAN plate reader (Series Infinite M200PRO). To determine the iron chelating capability of the bacterial supernatants the following formula was used: iron chelation 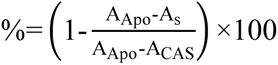.

A_Apo_: absorbance of uninoculated media + APO-CAS (positive control)

A_CAS_: absorbance of uninoculated media + CAS (negative control)

A_s_: absorbance of sample

### Bioinformatic analysis

Sequences of 35 *P. aeruginosa* PA01 TBDTs were obtained from Pseudomonas.com. Full length protein sequences were aligned using Clustalo (47). Individual domains were identified using HMMer3 (48) and Pfam domains Plug: PF07715.19 and TonB_dep_Rep: PF00593.28. Phylogenies were calculated using IQTree2 with the best model selected by ModelFinder (49). Phylogenies were visualised using iTOL (50). To analyse promoter regions, the upstream 500bp from each TBDT were accessed from Pseudomonas.com. Sequences were scanned against a combined database of prokaryote DNA motifs using SEA to identify enriched promoters against a control of shuffled input sequences. Sequences were then scanned with Fur (MX000013) and Zur (MX000374) binding site motifs. Hits was a p-value < 1×10^-3^ were considered significant. No significant hits were found for known copper regulatory motifs for any TBDTs.

---

**Fig. S1.**
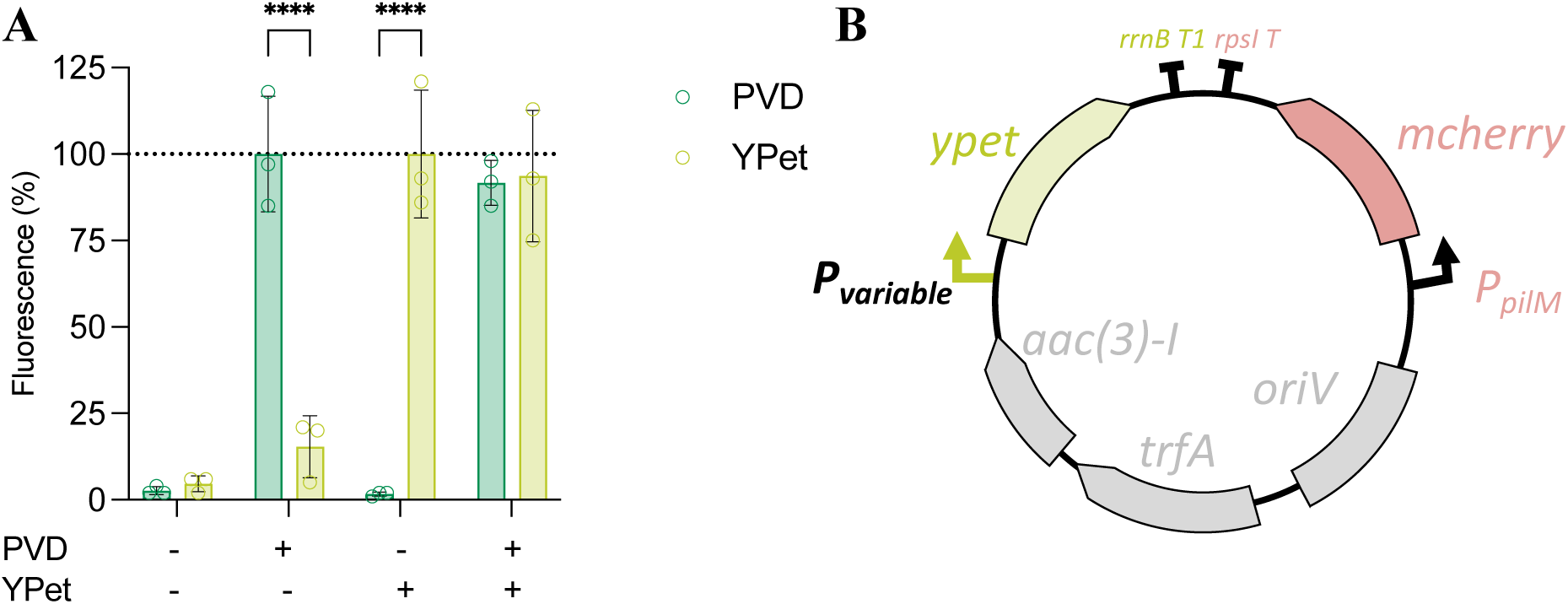
Transcriptional reporter plasmid for TBDT promoter activity measurement. (A) Fluorescence intensity of pyoverdine (ex: 400 nm em: 470 nm) and YPet (ex: 500 nm em: 540 nm). Fluorescence of a 20-hour culture of *P. aeruginosa* WT (PVD+) or *pchA pvdF cntL* mutant (PVD-), carrying pOPC-250 (P*btuB*) (YPet+) or not (YPet-) grown in CAA medium was measured on a Tecan plate reader. Error bars represent standard deviation from three independent measurements. (B) Schematic representation of reporter plasmids (variable promoter upstream of *ypet* fluorescent gene followed by *rrnB* terminator, for inducible YPet expression; P*_pilM_ mcherry* followed by *rpsI* terminator, for constitutive mCherry expression; *aac*(*3*)*-I*, conferring gentamicin resistance; *trfA*, replication initiator protein; *oriV*, origin of replication).

**Fig. S2.**
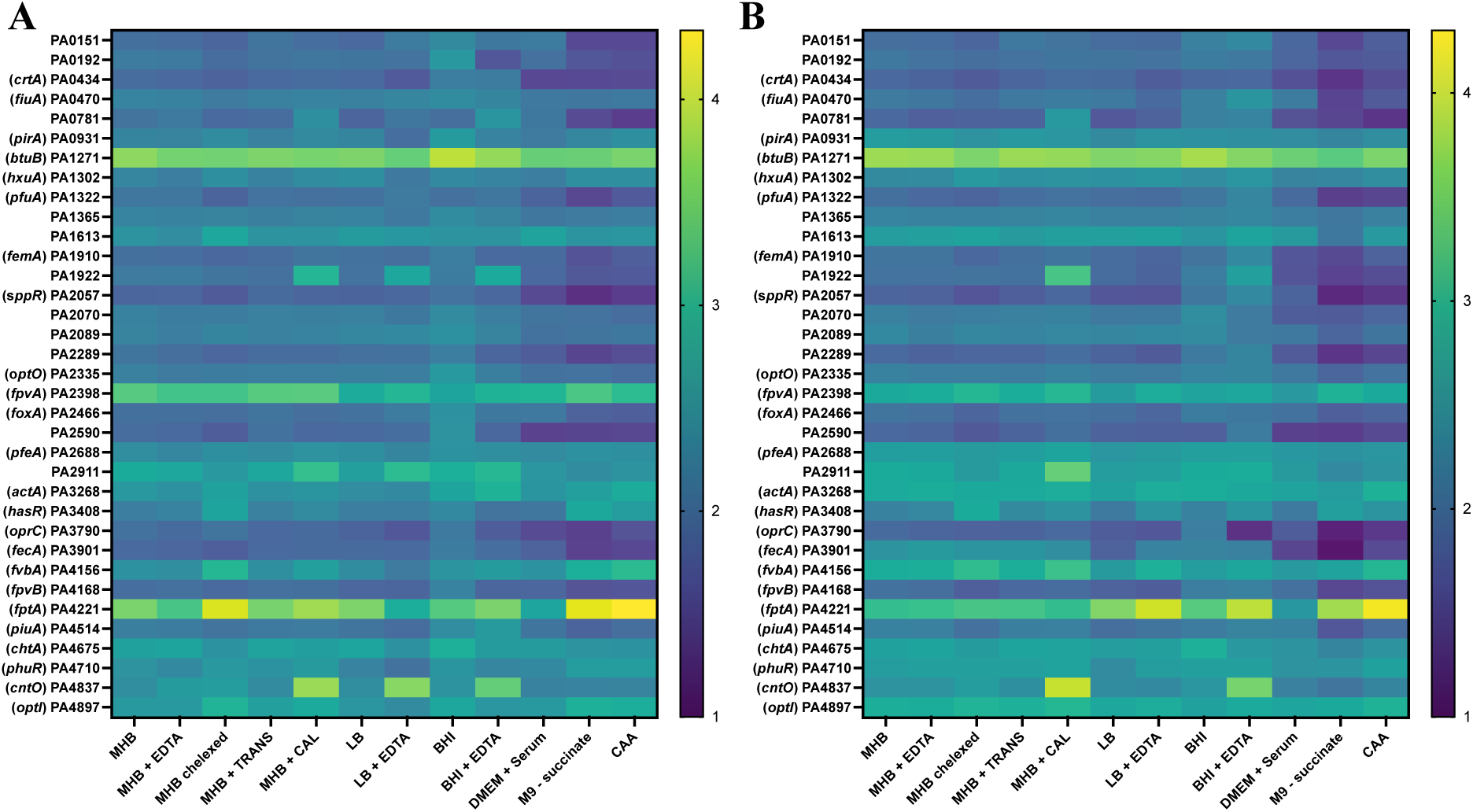
*P. aeruginosa* TBDT promoter activity in various growth condition. Log10 transformed values of *P. aeruginosa* PAO1 WT (A) and its isogenic *pchA pvdF cntL* mutant (B) TBDT promoter activity measured by fluorescence (YPet)/ optical density at 600 nm. The colour code goes from dark blue (least expression) over green (medium expression) to yellow (higher expression). Bacteria carrying TBDT reporter plasmids were grown media indicated on x-axis with supplementation of various additives for 20 h. MHB: + 300 µM of EDTA; + 30 µg/ml of apo-transferrin; + 30 µg/ml of calprotectin. LB: + 30 µM of EDTA. BHI: +100 µM of EDTA.

**Fig. S3.**
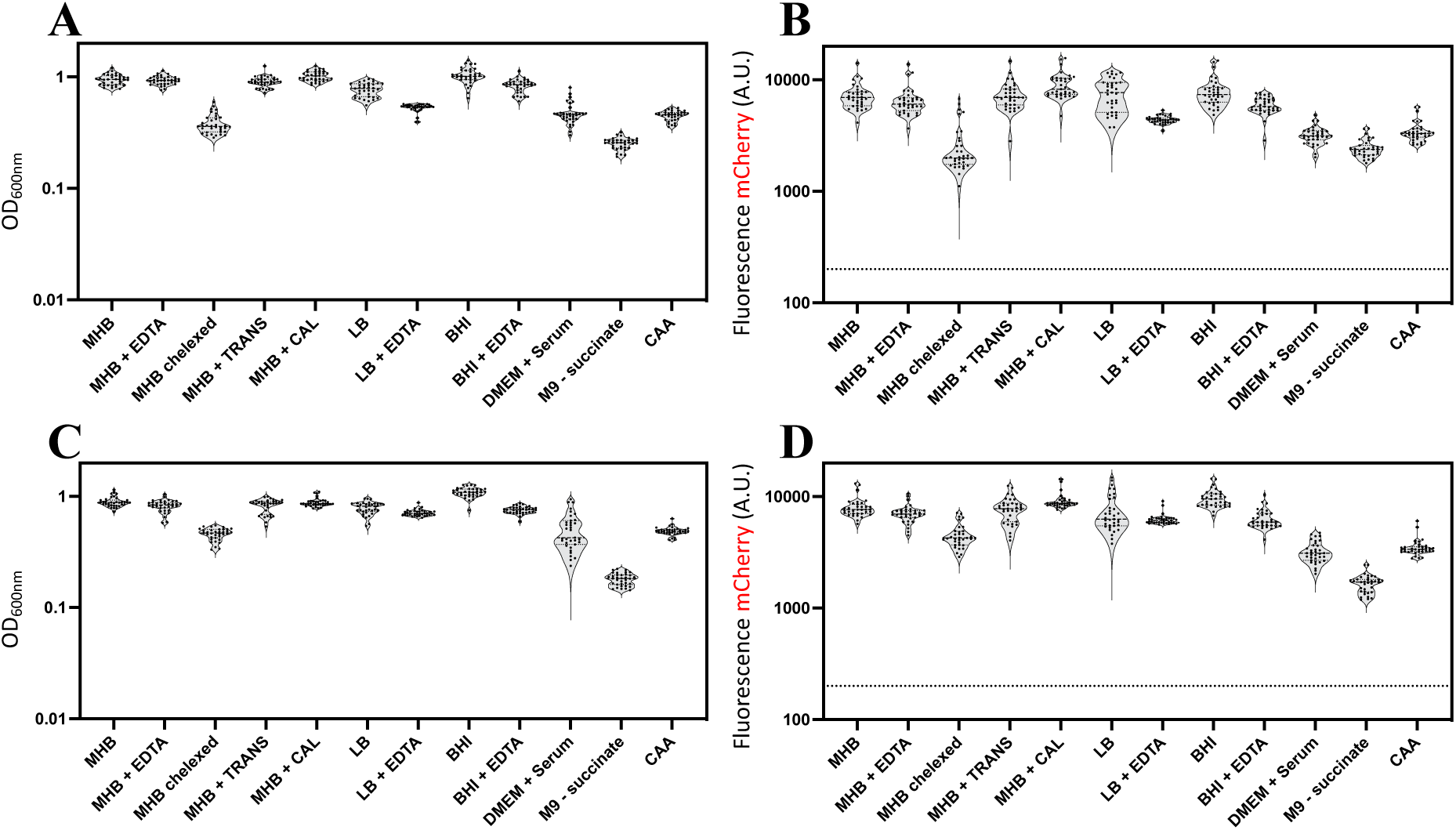
*P. aeruginosa* growth and mCherry fluorescence. (A) OD 600 nm values for *P. aeruginosa* WT strains carrying reporter constructs grown in various media for 20 h described in Fig. 1 and fig. S2. (B) mCherry values for *P. aeruginosa* WT strains carrying reporter constructs grown in various media for 20h described in Fig. 1 and fig. S2. (C) OD 600 nm values for *P. aeruginosa pchA pvdF cntL* strains carrying reporter constructs grown in various media for 20 h described in Fig. 1 and fig. S2. (D) mCherry values for *P. aeruginosa pchA pvdF cntL* strains carrying reporter constructs grown in various media for 20 h described in Fig. 1 and fig. S2.

**Fig. S4.**
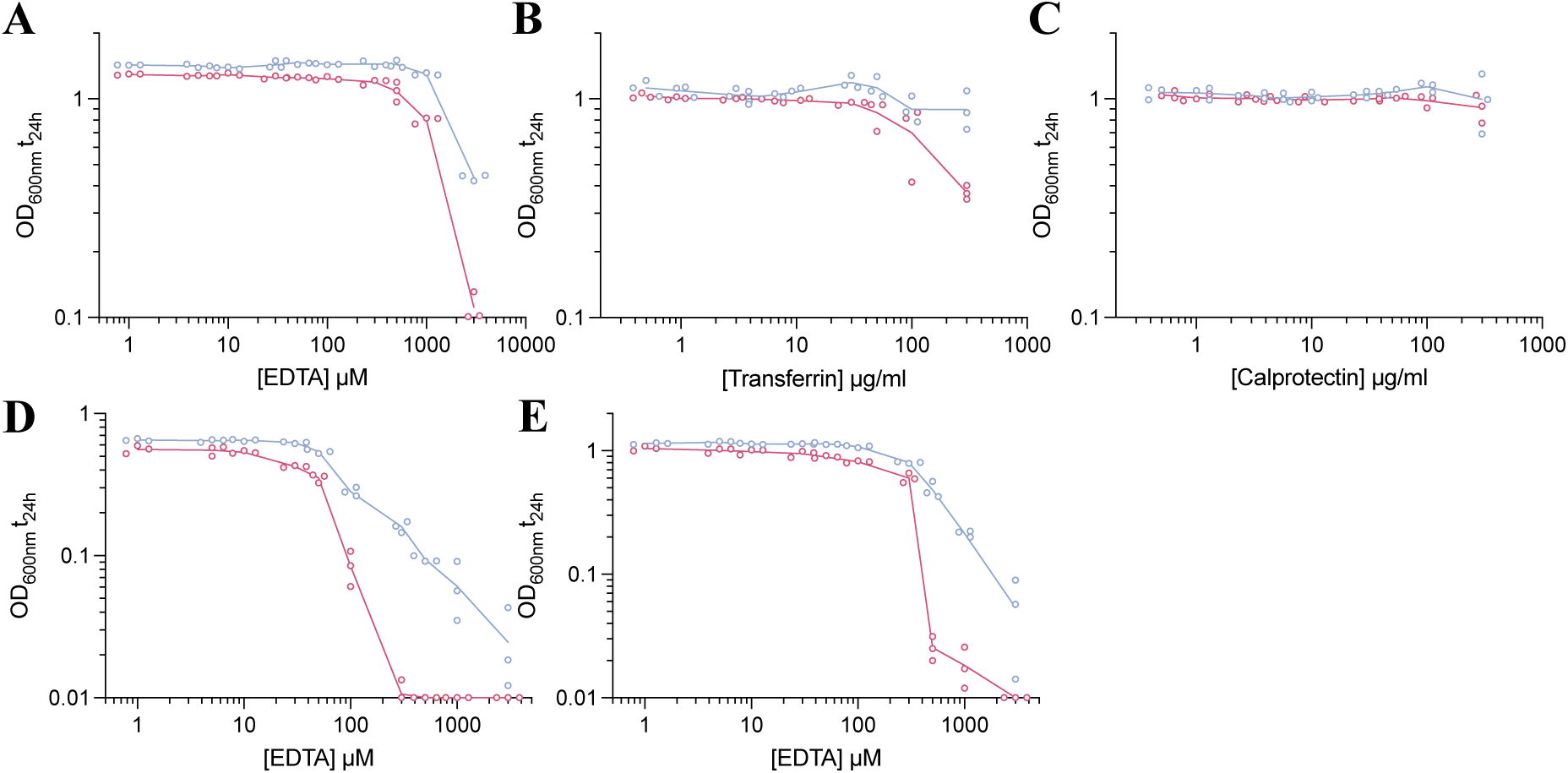
Growth inhibition experiment of *P. aeruginosa*. (A) OD 600 nm values of *P. aeruginosa* WT (blue) and *pchA pvdF cntL* (pink) grown in MHB with increasing concentration of EDTA. (B) OD 600 nm values of *P. aeruginosa* WT (blue) and *pchA pvdF cntL* (pink) grown in MHB with increasing concentration of apo-transferrin. (C) OD 600 nm values of *P. aeruginosa* WT (blue) and *pchA pvdF cntL* (pink) grown in MHB with increasing concentration of calprotectin. (D) OD 600 nm values of *P. aeruginosa* WT (blue) and *pchA pvdF cntL* (pink) grown in LB with increasing concentration of EDTA. (E) OD 600 nm values of *P. aeruginosa* WT (blue) and *pchA pvdF cntL* (pink) grown in BHI with increasing concentration of EDTA.

**Fig. S5.**
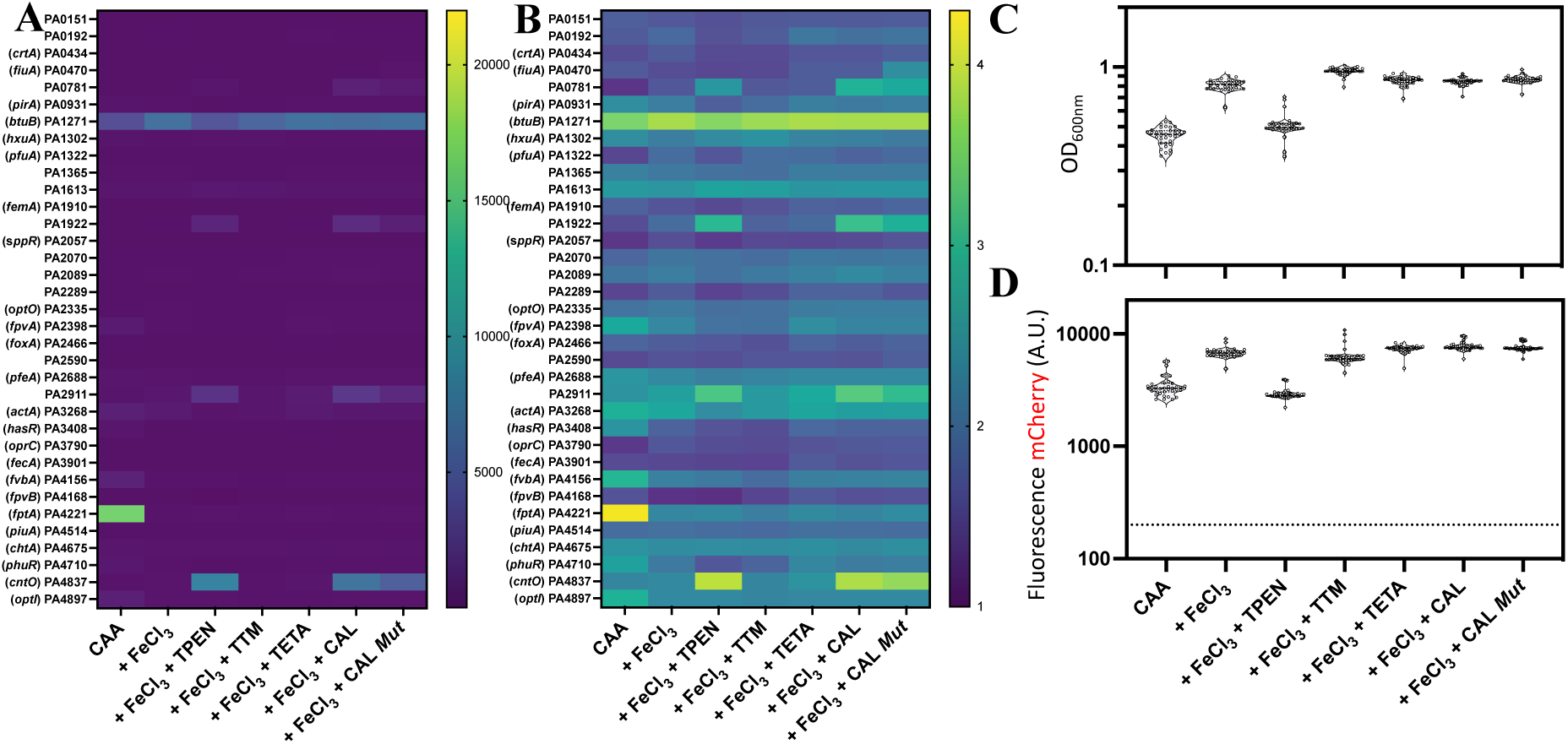
*P. aeruginosa* TBDT promoter activity is metal responsive. (A) Absolute values and (B) Log10 values of *P. aeruginosa* PAO1 *pchA pvdF cntL* mutant. TBDT promoter activity measured by fluorescence (YPet)/ optical density at 600 nm. Bacteria carrying TBDT reporter plasmids were grown in CAA medium at 37°C for 20 h, shaking supplemented with 10 µM of (FeCl_3_; TPEN; TTM, TETA) or 30 µg/ml of (CAL, CAL *Mut*). (C) OD 600 nm values for *P. aeruginosa* PAO1 *pchA pvdF cntL* mutant carrying reporter constructs grown in CAA described above and Fig. 2A. (B) mCherry values for *P. aeruginosa* PAO1 *pchA pvdF cntL* mutant carrying reporter constructs grown in CAA described above and Fig. 2A.

**Fig. S6.**
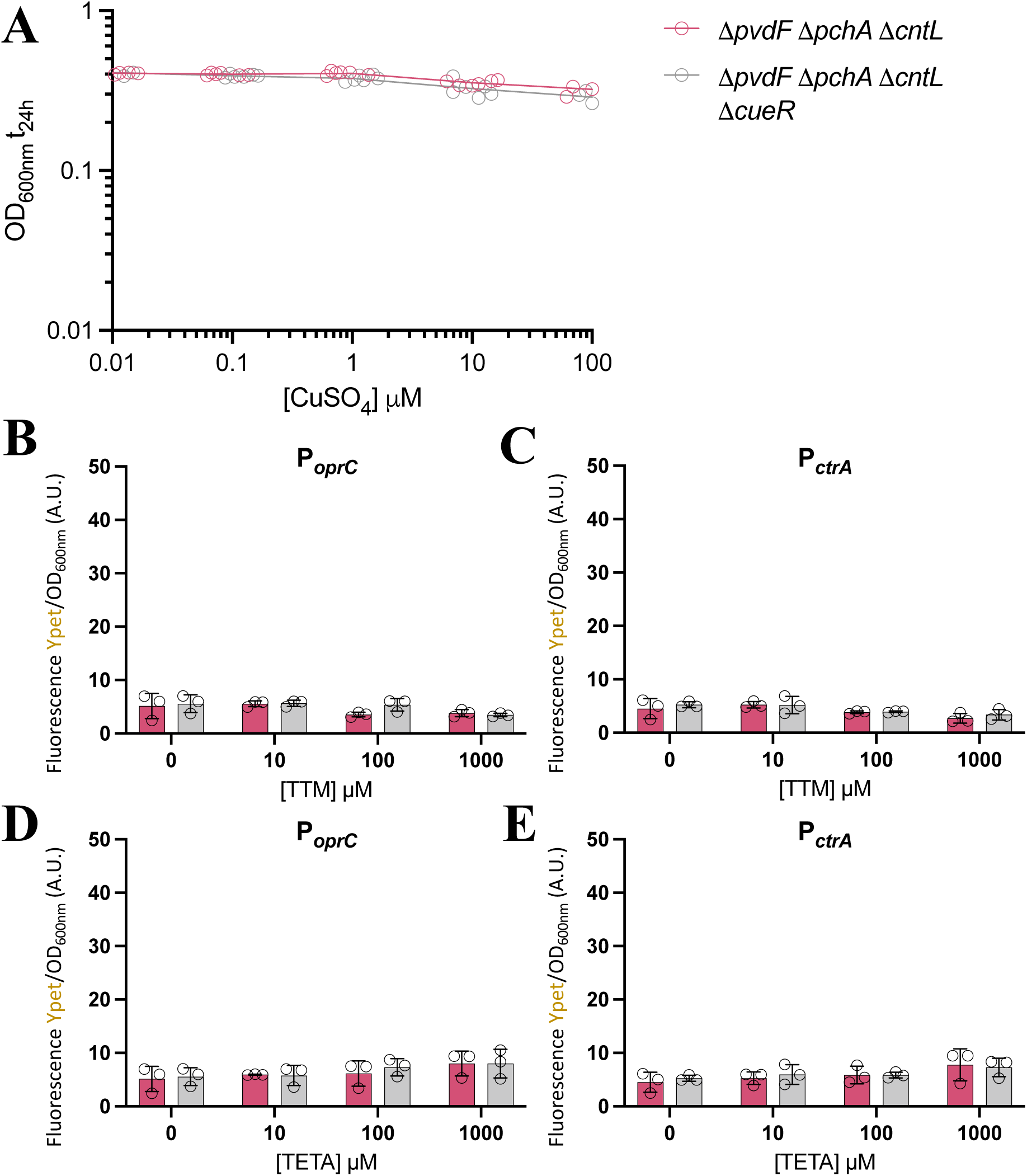
No measurable promoter activity during copper starvation. (A) OD 600 nm values of *P. aeruginosa pchA pvdF cntL* (pink) and *pchA pvdF cntL cueR* (grey) grown in CAA with increasing concentration of CuSO_4_. (B-E) Absolute values of *P. aeruginosa* PAO1 *pchA pvdF cntL* (pink) and *pchA pvdF cntL cueR* (grey) promoter activity. (B, D) P*oprC* and (C, E) P*PA0434* promoter activity was measured by fluorescence (YPet)/ optical density at 600 nm in increasing concentration of TTM (B, C) and TETA (D, E).

**Fig. S7.**
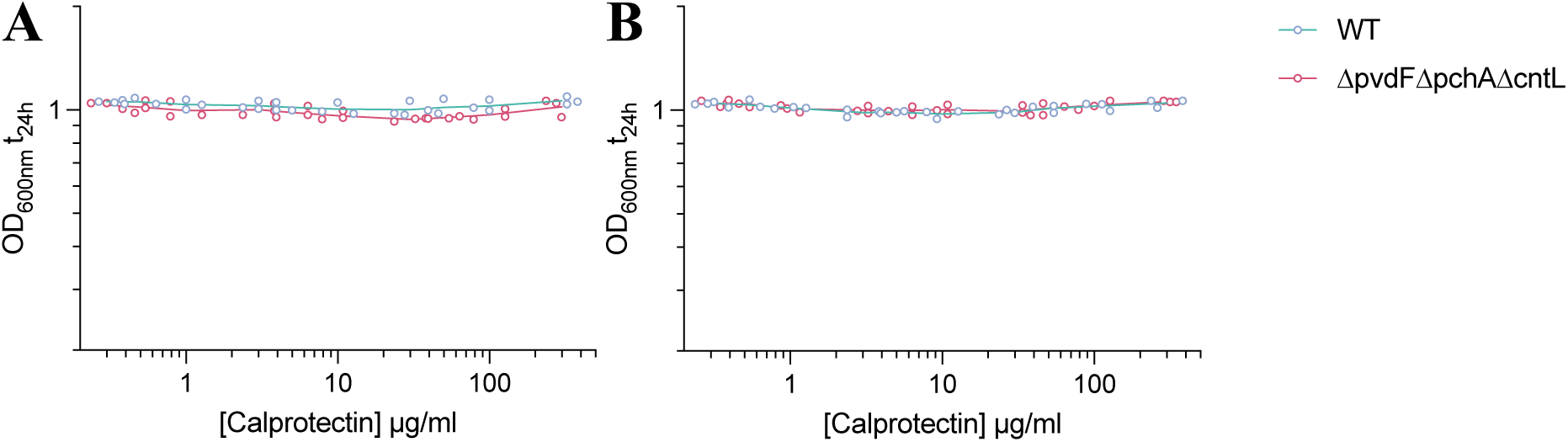
Growth inhibition experiment of *P. aeruginosa*. (A) OD 600 nm values of *P. aeruginosa* WT (blue) and *pchA pvdF cntL* (pink) grown in CAA supplemented with 10 µM of FeCl_3_ and increasing concentration of calprotectin. (B) OD 600 nm values of *P. aeruginosa* WT (blue) and *pchA pvdF cntL* (pink) grown in CAA supplemented with 10 µM of FeCl_3_ and increasing concentration of calprotectin *Mut* (H103N, H104N, and H105N).

**Fig. S8.**
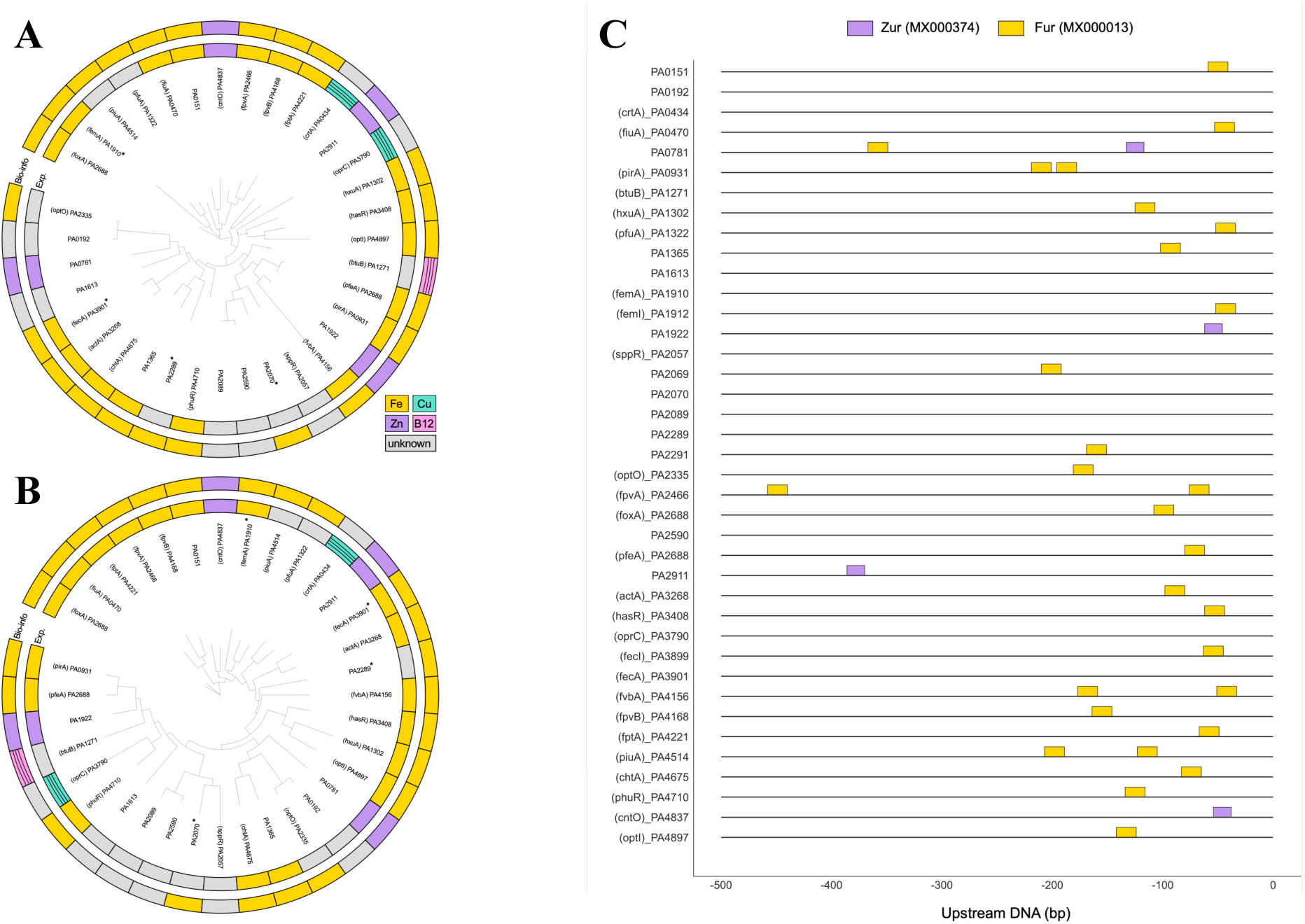
Bioinformatic analysis of *P. aeruginosa’s* 35 TBDTs. (A) Phylogeny of Plug domains (PF07715.19) from all TBDTs. (B) Phylogeny of regions within the TonB_dep_Rep domain (PF00593.28). Inner ring indicates experimental prediction of import function. Outer ring shows the prediction from bioinformatic analysis of promoter region. (C) Fur and Zur boxes identified in the upstream DNA of all 35 TBDT. All hits shown have a p-value < 0.001. In the upstream region of PA0781 where both Zur and Fur were identified the hit with the strongest p-value was chosen.

**Fig. S9.**
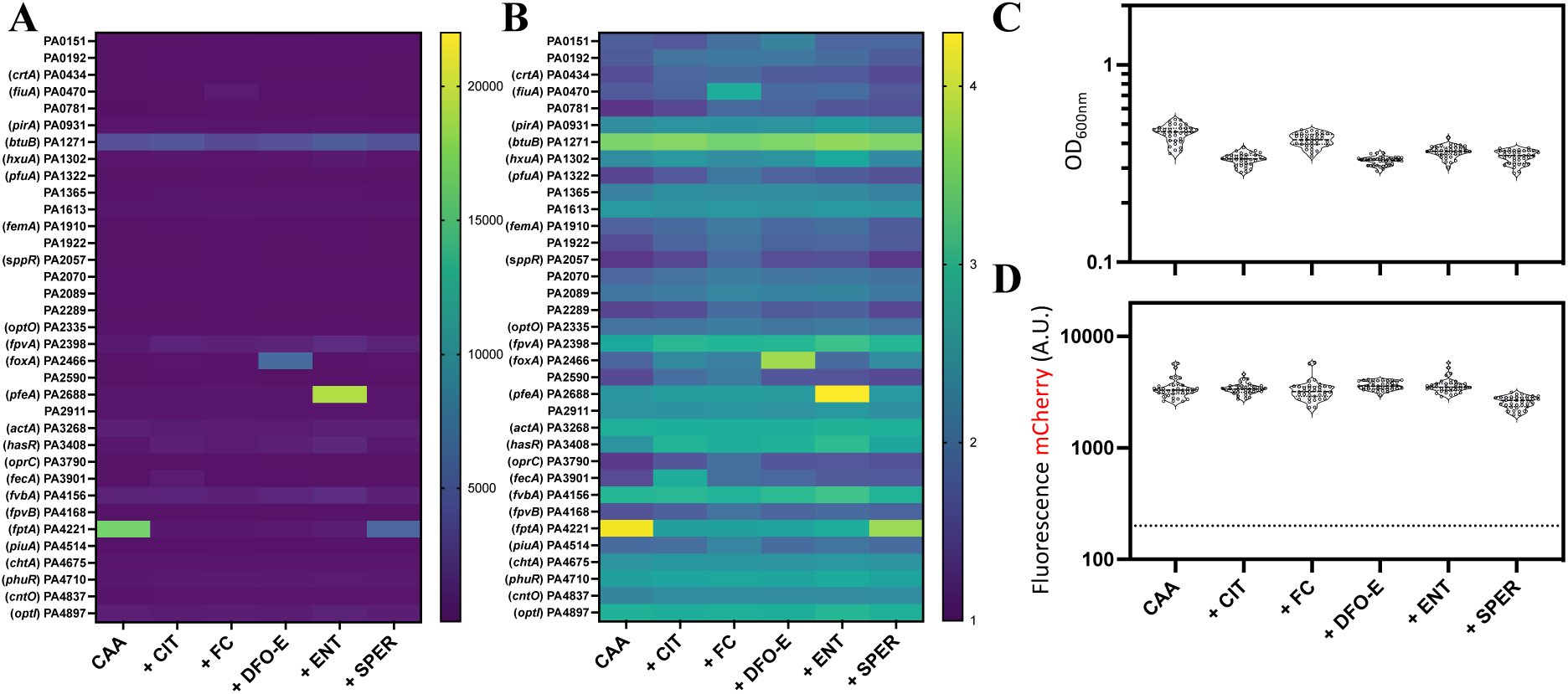
Reporter constructs detect single siderophores. (A) Absolute values and (B) Log10 values of *P. aeruginosa* PAO1 *pchA pvdF cntL* mutant. TBDT promoter activity measured by fluorescence (YPet)/ optical density at 600 nm. Bacteria carrying TBDT reporter plasmids were grown in CAA medium at 37°C for 20h, shaking supplemented with 1 mM of CIT – citrate, or 10 µM of (FC - ferrichrome; DFO-E – desferroxamine E; ENT – enterobactin; SPER – spermidine). (C) OD 600 nm values for *P. aeruginosa* PAO1 *pchA pvdF cntL* mutant carrying reporter constructs grown in CAA described above and Fig. 3A. (D) mCherry values for *P. aeruginosa* PAO1 *pchA pvdF cntL* mutant carrying reporter constructs grown in CAA described above and Fig. 3A.

**Fig. S10.**
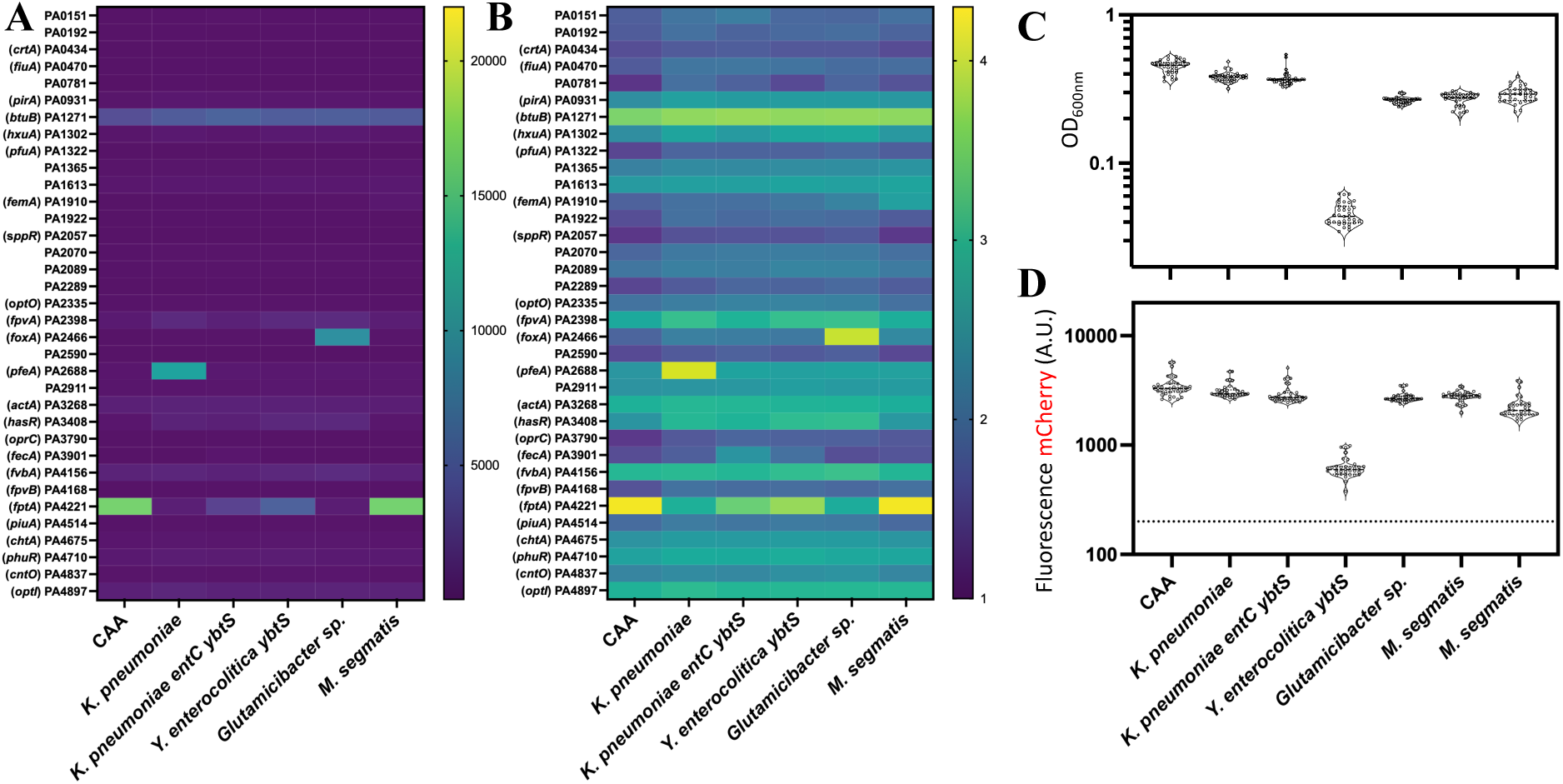
Reporter constructs detect siderophores in bacterial supernatants. (A) Absolute values and (B) Log10 values of *P. aeruginosa* PAO1 *pchA pvdF cntL* mutant. TBDT promoter activity measured by fluorescence (YPet)/ optical density at 600 nm. Bacteria carrying TBDT reporter plasmids were grown in CAA medium at 37°C for 20 h, shaking supplemented with 1/1 bacterial supernatants. (C) OD 600 nm values for *P. aeruginosa* PAO1 *pchA pvdF cntL* mutant carrying reporter constructs grown in CAA described above and Fig. 3B. (D) mCherry values for *P. aeruginosa* PAO1 *pchA pvdF cntL* mutant carrying reporter constructs grown in CAA described above and Fig. 3B.

**Fig. S11.**
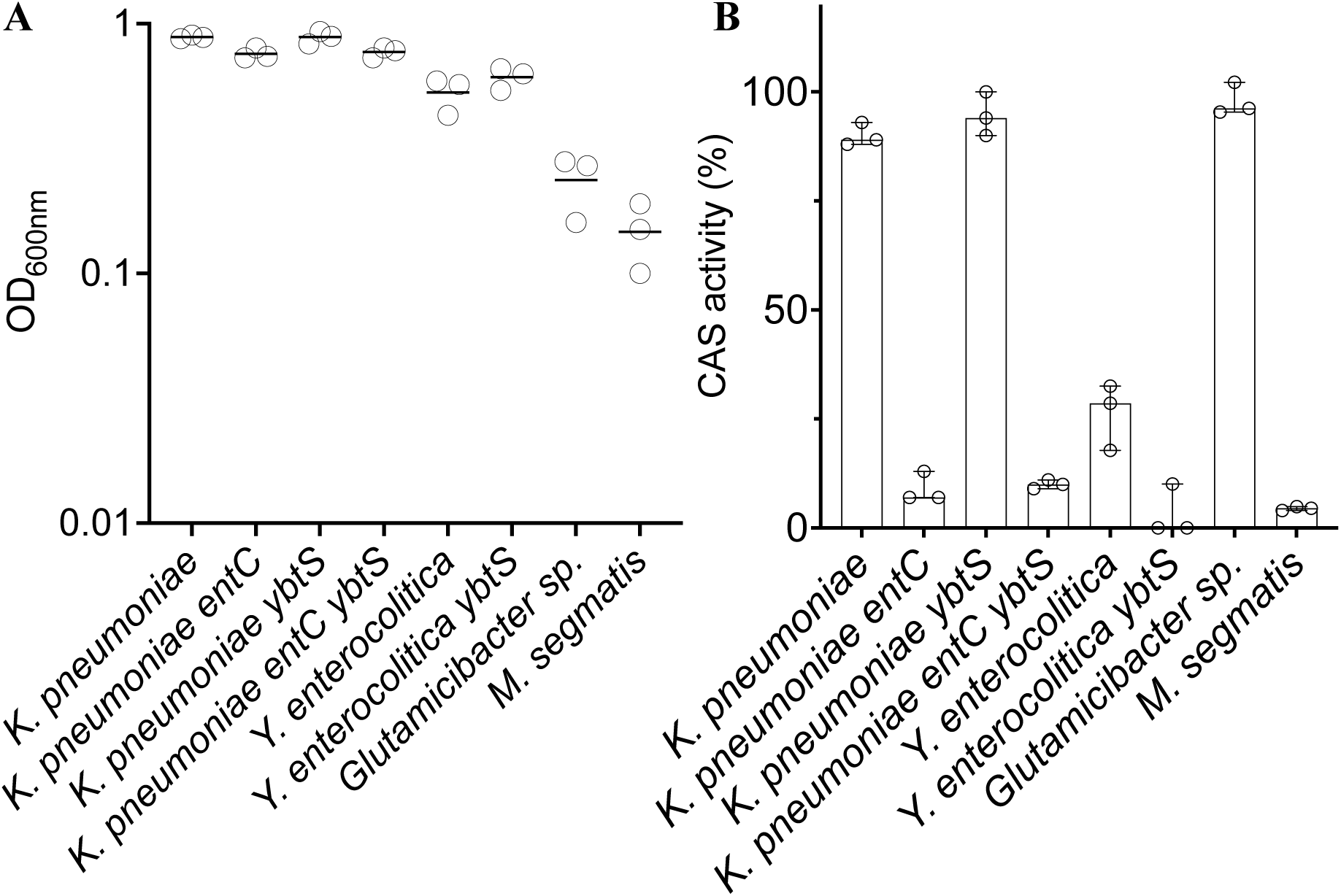
Growth characteristics and siderophore production. (A) OD 600 nm values of bacteria grown in iron restricted medium. *K. pneumoniae* and *M. smegmatis* grown in low phosphate M9; and *Y. enterocolitica* and *Glutamicibacter sp*. were grown in CAA medium. (B) Culture supernatant were used to measure siderophore production using the CAS assay, described in Method part.

**Fig. S12.**
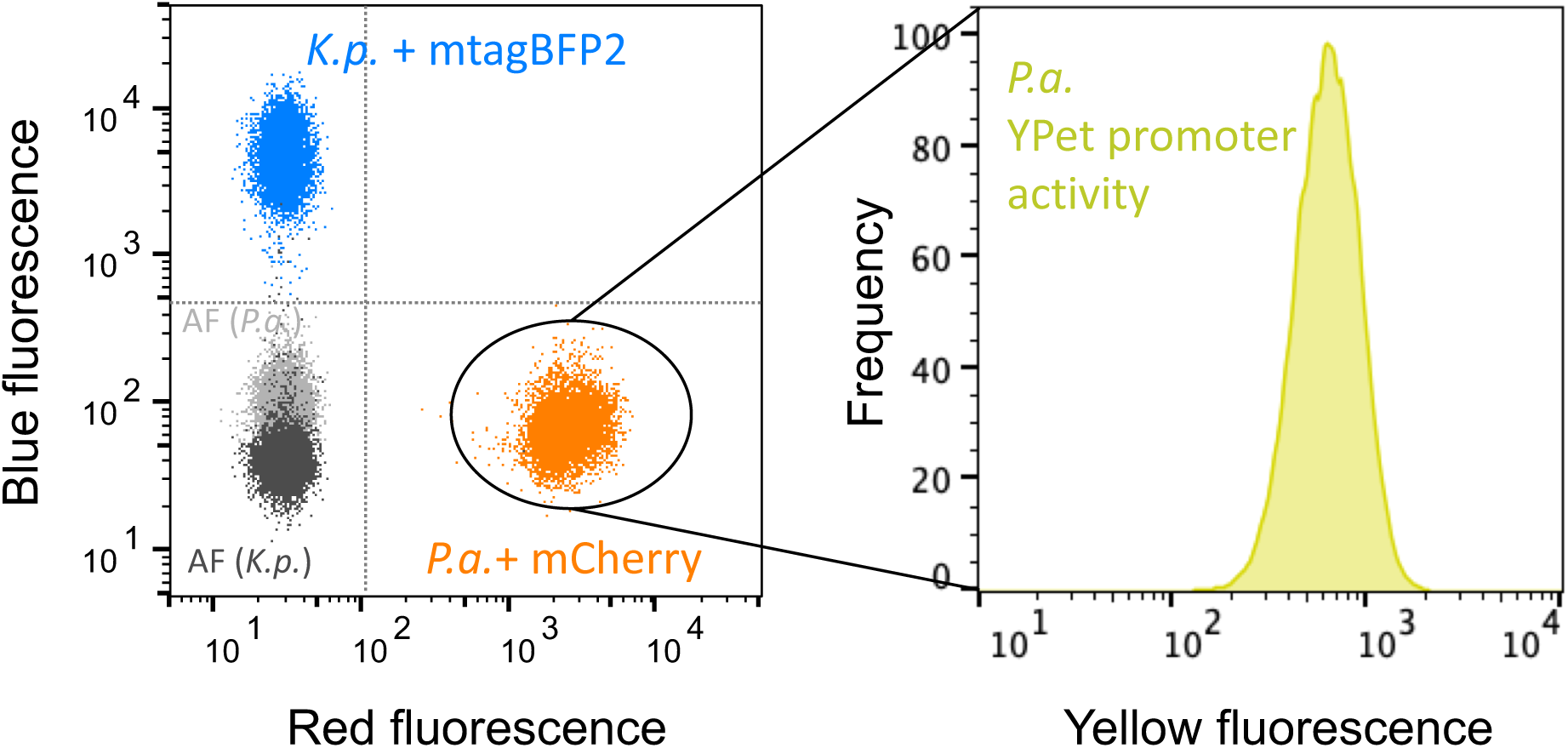
Gating strategy. Flow cytometry data are presented with each dot representing an individual particle. *Pseudomonas* strains express mCherry constitutively, enabling detection through red fluorescence, while *Klebsiella* strains express mtagBFP2, which provides blue fluorescence for discrimination in co-cultures. The yellow fluorescence of YPet in *Pseudomonas* is assessed as an indicator of gene expression. Autofluorescence for both species is shown in gray and black. Background noise was minimized using forward and side scatter.

## Notes

### Competing Interest Statement

The authors have declared no competing interest.

